# Discovery and Metabolic Origin of 4,4′-Dihydroxy-3,3′,5,5′-Tetrachlorobenzophenone from a *Burkholderia oklahomensis* Clinical Isolate

**DOI:** 10.1101/2025.05.18.654707

**Authors:** Yousef Dashti, Guy J. Clarkson, Eshwar Mahenthiralingam, Gregory L Challis

## Abstract

*Burkholderia oklahomensis* LMG 23618^T^ is a *Burkholderia pseudomallei*-like bacterium originally isolated in 1973 from a wound infection caused by a farming accident in Oklahoma. Metabolic profiling of an organic extract from cultures of *B. oklahomensis* LMG 23618^T^ using UHPLC-ESI-Q-ToF-MS led to identification of three known metabolites, betulinan A, yersiniabactin and ulbactin B, in addition to a novel polycholorinated compound. Mass-directed purification enabled isolation of the novel specialized metabolite, which was shown by X-ray crystallography and NMR spectroscopic analysis to be 4,4′-dihydroxy-3,3′,5,5′-tetrachlorobenzophenone. Feeding experiments with stable isotope-labelled precursors established that the carbon skeleton of this unusual metabolite derives from two molecules of tyrosine. This led us to propose a plausible biosynthetic pathway via decarboxylative condensation of 3, 5-dichloro-4-hydroxybenzoic acid with its coenzyme A thioester derivative. The absolute configuration of ulbactin B was also established as 4′R, 3″S, 7″S, 8″R using X-ray crystallography and NMR spectroscopy.

## Introduction

Specialized metabolites produced by microbes are a rich source of novel molecular scaffolds with potent biological activities. Many clinically approved medicines are, or have origins in microbial specialized metabolites.^1^ Traditionally, drug discovery programs have focused on Actinomycetes, but advances in genome sequencing and bioinformatics have revealed that underexplored bacterial taxa also harbor great potential for the production of novel specialized metabolites.^2-4^

The genus *Burkholderia* sensu lato encompasses ecologically diverse Gram-negative bacteria that inhabit a wide range of terrestrial and aquatic environments. These bacteria can be free-living or exist in association with a variety of hosts, including humans, animals, plants, and fungi.^5, 6^ Their interactions with host organisms range from harmful, such as opportunistic infections caused by members of the *Burkholderia cepacia* complex (Bcc) in cystic fibrosis patients,^7^ to beneficial, promoting plant growth^8, 9^ or acting as biopesticides by protecting crops from pathogens.^10, 11^

*Burkholderia* species are prolific producers of specialised metabolites, and genomic data suggest they harbour a vast untapped potential to assemble novel compounds with potential applications in medicine and agriculture.^4, 11-13^ In our ongoing search for novel natural products from human pathogenic *Burkholderia* species,^13-17^ we have identified the novel specialised metabolite 4, 4′-dihydroxy-3, 3′, 5, 5′-tetrachlorobenzophenone (**1)** from *B. oklahomensis* LMG 23618^T^, along with known metabolites betulinan A (**2**), yersiniabactin (**3**), and ulbactin B (**4**) (Figure 1).^18^ Incorporation experiments with stable isotope-labelled precursors demonstrate that the carbon skeleton of **1** derives from two molecules of tyrosine, leading us to propose a plausible biosynthetic pathway to this unusual metabolite.

**Figure 1.**
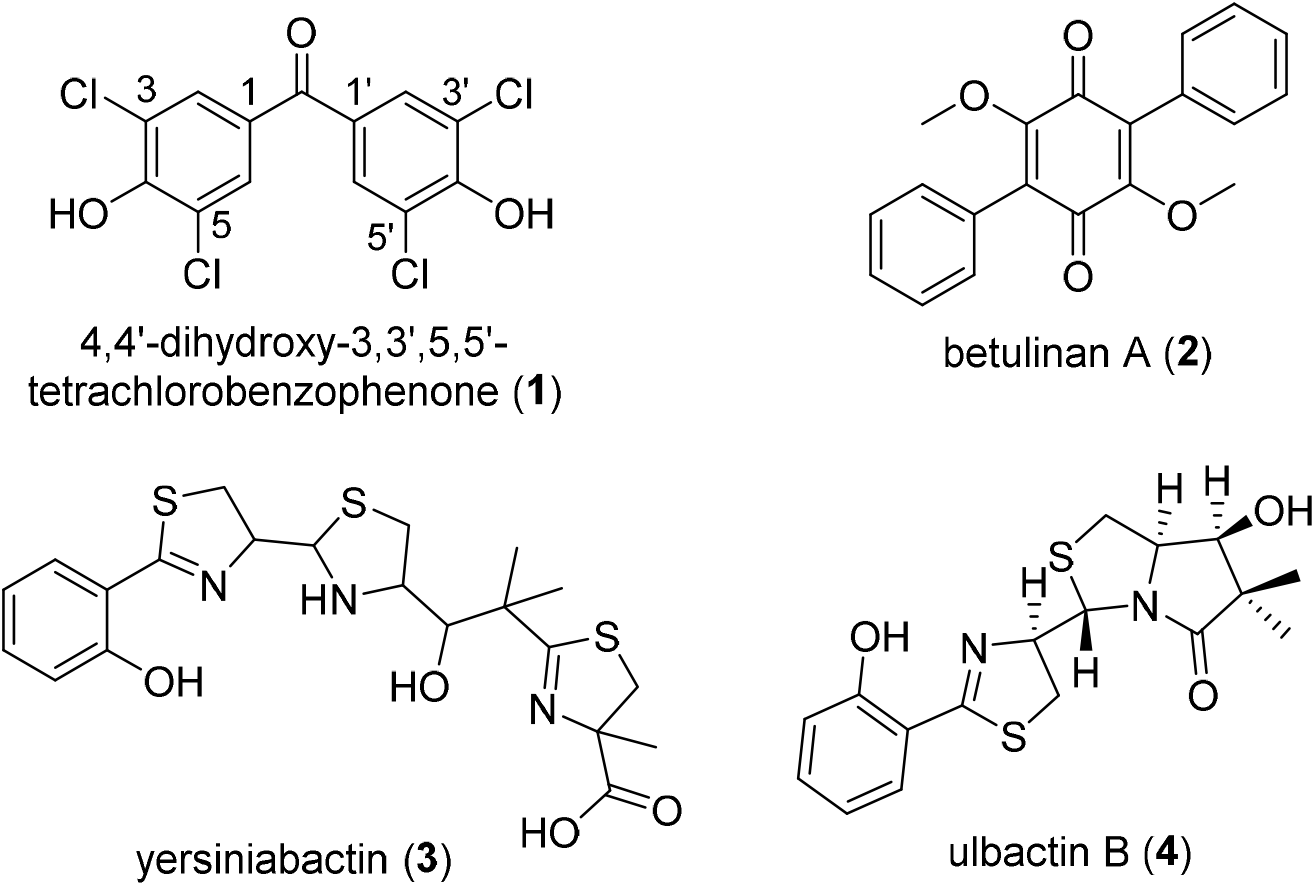
Structures of 4,4′-dihydroxy-3,3′,5,5′-tetrachlorobenzophenone (**1**), betulinan A (**2**), yersiniabactin (**3**), and ulbactin B (**4**) isolated from *B. oklahomensis* LMG 23618^T^.

## Results and discussion

The specialized metabolite profile of an ethyl acetate extract from *B. oklahomensis* LMG 23618^T^, cultured on Basal Salts Medium (BSM) supplemented with glycerol as the carbon source was analysed by UHPLC-ESI-Q-ToF-MS. Metabolites with molecular formulae corresponding to three known metabolites (**2, 3**, and **4**) were identified in the extract, alongside a novel natural product with the molecular formula C_13_H_6_Cl_4_O (calculated *m/z* = 350.9144 for [M+H]^+^; measured *m/z* = 350.9147). The mass spectrum of the novel metabolite exhibited the expected isotope distribution for a tetracholorinated compound with signals of the expected intensity at *m/z* = 350.9147, 352.9114, 354.9089, and 356.9059, corresponding to the [M+H]^+^ ions for the ^35^Cl_4_, ^35^Cl_3_^37^Cl, ^35^Cl_2_^37^Cl_2_ and ^35^Cl^37^Cl_3_ isotopes. Mass-directed purification enabled the isolation of the compound, which was analysed using ^1^H, ^13^C, COSY, HSQC, and HMBC NMR experiments. The ^1^H NMR spectrum had a single resonance at *δ*_*H*_ 7.67 ppm. In the ^13^C NMR spectrum, four quaternary carbon signals were observed at *δ*_*C*_ 122.1, 128.8, 153.6, and 189.2 ppm, along with one sp^2^ methine carbon at *δ*_*C*_ 130.1 ppm. The sole proton signal showed an HSQC correlation to the methine carbon, and exhibited ^*2*^*J* and ^*3*^*J* HMBC correlations to the four quaternary carbons.

While the MS and NMR data were consistent with a symmetrical tetrachlorinated architecture, they did not allow unambiguous assignment of the structure. We therefore grew crystals for analysis by X-ray diffraction. Suitable single crystals were obtained by slow diffusion of hexane into an ethyl acetate solution of **1**. The resulting X-ray crystal structure showed that **1** is 4,4′-dihydroxy-3,3′,5,5′-tetrachlorobenzophenone (Figure 2). The symmetrical nature of this unusual tetrachlorinated metabolite explains the presence of only a single resonance in the ^1^H NMR spectrum and five distinct signals in the ^13^C spectrum.

**Figure 2.**
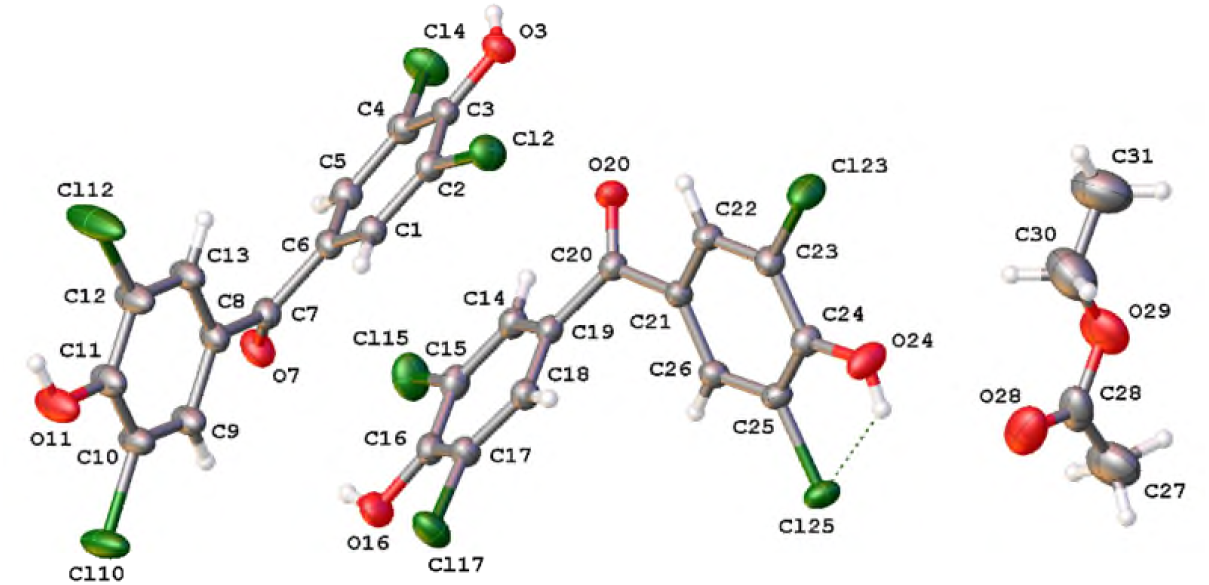
Asymmetric unit of 4,4′-dihydroxy-3,3′,5,5′-tetrachlorobenzophenone (**1**) crystals with atom labelling and thermal ellipsoids drawn at 50% probability level. The molecule of ethyl acetate was refined at 50% occupancy.

To elucidate the metabolic origin of **1**, feeding experiments were conducted using 1 mM of uniformly ^13^C-labelled tyrosine and acetic acid, each supplemented separately into the culture medium. *B. oklahomensis* LMG 23618^T^ was grown in BSM for three days, after which extracts were analysed by UHPLC-ESI-Q-ToF-MS. No incorporation of ^13^C from acetic acid was detected. However, a high level of ^13^C incorporation from tyrosine was observed for all carbons in **1** (Figures 3a and 3b), confirming that its carbon skeleton derives from two molecules of tyrosine.

**Figure 3.**
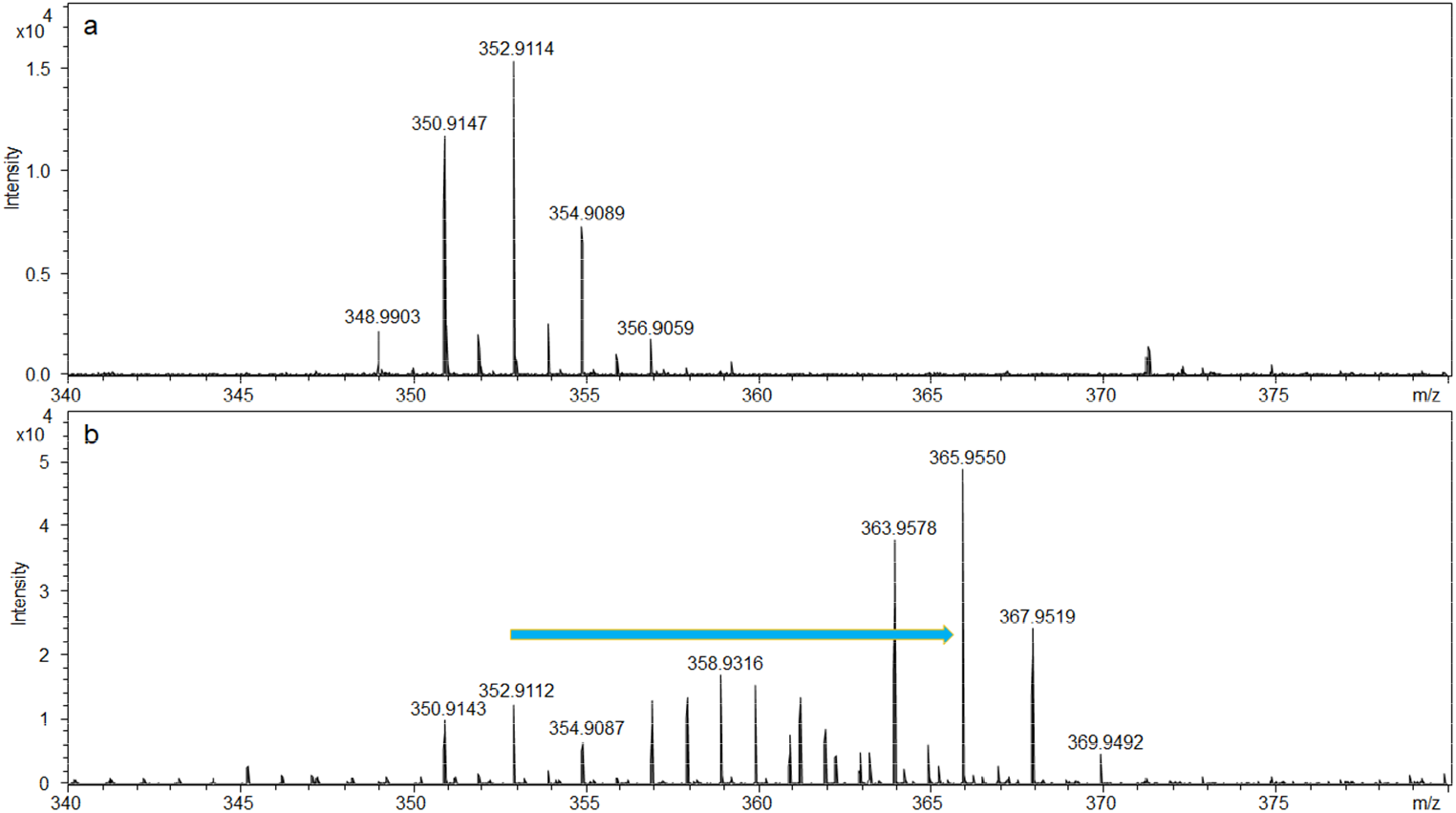
Incorporation of uniformly ^13^C-labelled tyrosine into 4,4′-dihydroxy-3,3′,5,5′-tetrachlorobenzophenone (**1**). **a**) Mass spectrum of **1** in the extract from *B. oklahomensis* cultured in BSM medium. **b**) Mass spectrum of **1** in the extract from *B. oklahomensis* cultured in BSM medium supplemented with uniformly ^13^C-labelled tyrosine. A shift of 13 a.m.u. in the latter indicates all carbon atoms of **1** derive from tyrosine.

Given that L-tyrosine is known to be degraded to *p*-hydroxybenzoic acid (*p*HBA),^19, 20^ we propose **1** is biosynthesised as follows (Figure 4). Ammonia is eliminated from L-tyrosine by tyrosine ammonia lyase (TAL) yielding *p*-coumaric acid (*p*CA), which is subsequently converted to the corresponding coenzyme (CoA) thioester by feruloyl-CoA synthetase (Fcs). The activated intermediate undergoes hydration and retro-Aldol cleavage between the *α* and *β* carbons, catalysed by enoyl-CoA hydratase (Ech), resulting in *p*-hydroxybenzaldehyde (*p*HBAL) and acetyl-CoA. Oxidation of *p*HBAL by vanillin dehydrogenase (Vdh) yields *p*HBA, which is dichlorinated at both *meta* carbons by an FADH_2_-dependent chlorinase. One molecule of *m, m*-dichloro-*p-*hydroxybenzoic acid (*mm*D*p*HBA) is converted to the corresponding CoA thioester by an acyl-CoA synthetase, enabling it to acylate the *ipso* position of the phenol in a second molecule of *mm*D*p*HBA. Decarboxylation of the resulting intermediate restores aromaticity resulting in the formation of **1**. It is currently unclear what type of enzyme might catalyse the final acylation-decarboxylation sequence.

**Figure 4.**
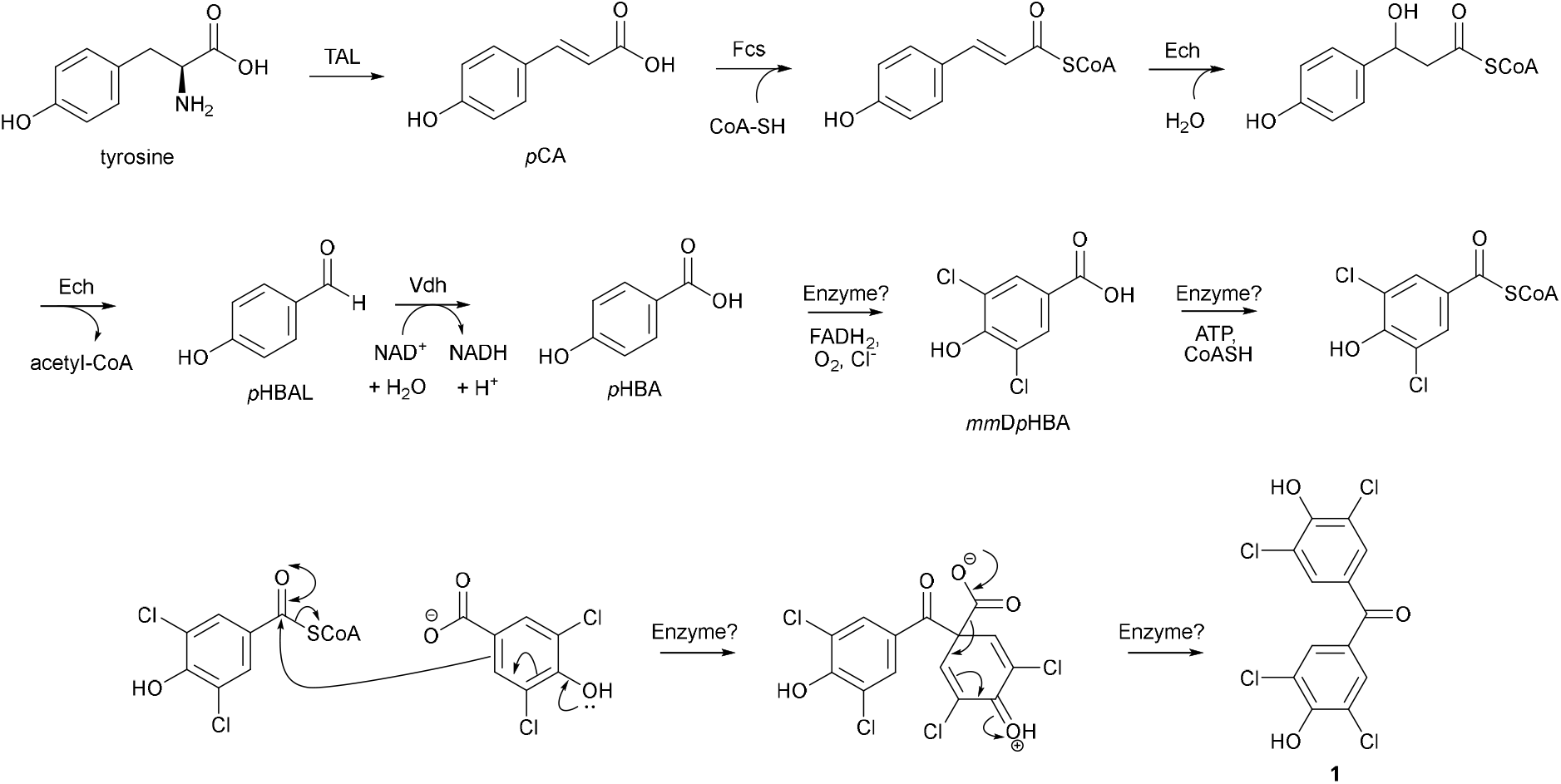
Proposed biosynthesis of 4,4′-dihydroxy-3,3′,5,5′-tetrachlorobenzophenone (**1**) from tyrosine. The enzymes that catalyse the chlorination, acyl-CoA formation, and acylation-decarboxyaltion reactions remain to be identified.

Yersiniabactin (**3**) and ulbactin B (**4**) are structurally related metabolites identified in the extract of *B. oklahomensis*. Yersiniabactin is a siderophore previously reported to be produced by several pathogenic bacteria, including *Yersinia pestis, Y. pseudotuberculosis, Y. enterocolitica, E. coli*, and *Salmonella enterica*.^21-24^ Like other siderophores, yersiniabactin plays a crucial role in bacterial pathogenicity by facilitating iron acquisition.^25^ The structurally related siderophore pyochelin has been identified in other pathogenic *Burkholderia* species.^26^ To our knowledge this is the first report of yersiniabactin production in *Burkholderia*.

Since the initial discovery of ulbactin B from *Vibrio* sp. B-93 in 1996,^27^ its absolute configuration has remained unassigned. Using the same crystallization method employed for **1**, we obtained crystals of ulbactin B suitable for X-ray diffraction. This enabled its absolute configuration to be assigned as 4′R, 3′′S, 7′′S, 8′′R. This stereochemistry is supported by NOESY correlations between H-10” and H-7”, as well as between H-7” and H-8” (Figure 5).

**Figure 5.**
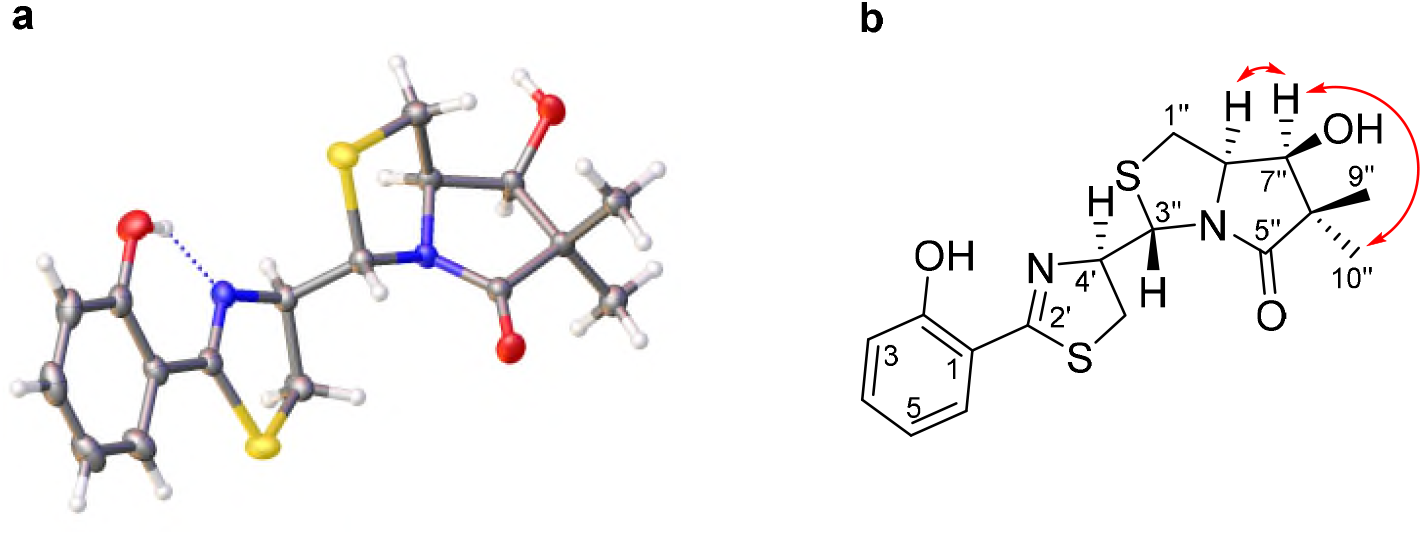
**a**) Solid-state molecular structure of ulbactin B (**4**) with absolute stereochemistry assigned as 4′R, 3′′S, 7′′S, 8′′R based on single-crystal X-ray diffraction analysis. **b)** Key NOESY correlations supporting the stereochemical assignment.

## Conclusion

This study reports the discovery and metabolic origin of the novel polychlorinated metabolite 4,4′-dihydroxy-3,3′,5,5′-tetrachlorobenzophenone from the clinical isolate *B. oklahomensis* LMG 23618^T^, along with three known metabolites betulinan A, yersiniabactin, and ulbactin B. Stable isotope feeding experiments revealed that the carbon framework of this unusual benzophenone is derived from two molecules of tyrosine, leading us to propose a plausible biosynthetic pathway involving decarboxylative condensation of halogenated 4-hydroxybenzoic acid derivatives. The absolute stereochemistry of ulbactin B was also determined for the first time using X-ray crystallography.

*B. oklahomensis* is closely related to *B. pseudomallei*, the causative agent of melioidosis, which a recent outbreak in northern Australia has shown leads to mortality in 20% of those infected. *B. oklahomensis* has been reported to be much less virulent than *B. pseudomallei*.^28^ However, our finding that it produces yersiniabactin, a siderophore that has been shown play an important role in the virulence of other bacteria, such as the plague pathogen *Yersinia pestis*,^29^ warrants further investigation. Interestingly, *B. pseudomallei* does not rely on yersiniabactin for iron acquisition, instead employing pyochelin and malleobactin, which appear not to play a critical role on virulence.^30^

## Materials and Methods

### General Experimental Procedures

UHPLC-ESI-Q-ToF-MS analyses were performed using a Dionex UltiMate 3000 UHPLC system coupled to a Bruker MaXis IMPACT mass spectrometer via a Zorbax Eclipse Plus C18 column (100 × 2.1 mm, 1.8 μm). The mobile phases consist of water and acetonitrile, both containing 0.1% formic acid. Chromatographic separation was achieved using a gradient elution from 5% to 100% acetonitrile over 30 minutes at a flow rate of 0.2 mL/min. The mass spectrometer was operated in positive ion mode with a scan range of *m/z* 50-3000. A 1 mM sodium formate solution was used for calibration, introduced via a 20 μL loop injection at the start of each run. NMR spectra were recorded on a Bruker 600 MHz spectrometer using DMSO-*d*_*6*_ as the solvent. ^1^H and ^13^C NMR chemical shifts were referenced to the DMSO-*d*_*6*_ solvent signals at *Δ*_H_ 2.50 and *Δ*_C_ 39.51, respectively. Single crystal X-ray diffraction data were collected on either a Rigaku Xcalibur Gemini diffractometer equipped with a Ruby CCD area detector or a Rigaku Oxford Diffraction SuperNova diffractometer with a dual source (Cu at zero) system and an AtlasS2 CCD area detector.

### Production and Purification of Specialized Metabolites from *B. oklahomensis*

*B. oklahomensis* LMG 23618^T 18^ (strain BCC1605 in the Cardiff University Collection^7^) was cultured on 1 L of solid Basal Salts Medium (BSM) supplemented with glycerol as the carbon source^31^ for 3 days at 30 °C. After incubation, the agar was chopped into small blocks and extracted twice with EtOAc. The combined EtOAc extracts were dried using a rotary evaporator, then resuspended on 1 mL MeOH and pre-adsorbed onto C18-bonded silica. The dried mixture was packed into a stainless steel HPLC guard cartridge (10 × 30 mm) and connected to a semi-preparative reverse-phase C18 Betasil column (21.2 × 150 mm). Purification was carried out using initial isocratic elution with 5% acetonitrile for 5 minutes, followed by a linear gradient from 5% to 100% acetonitrile over 45 minutes, and a final isocratic hold at 100% acetonitrile for 10 min. The fellow rate was maintained at 9 mL/min. Fractions were collected every minute for a total of 60 minutes. Ulbactin B was eluted in fraction 36, 4,4′-dihydroxy-3,3′,5,5′-tetrachlorobenzophenone was obtain pure in fractions 46, and betulinan A was eluted in fraction 47. The structure of the know compounds were confirmed by comparing the molecular formulas obtained via HR-MS and ^1^H NMR, and where necessary, 2D NMR, with published data.

4,4′-dihydroxy-3,3′,5,5′-tetrachlorobenzophenone (**1**): (1.3 mg); ^1^H NMR (600 MHz, DMSO-*d*_*6*_): *δ*_*H*_ 7.67; ^13^C NMR (150 MHz, DMSO-*d*_*6*_): *Δ*_C_,122.1 (C3, C5, C3′, C5′), 128.8 (C1, C1′), 130.1 (C2, C6, C2′, C6′), 153.6 (C4, C4′), 189.2 (C=O); HRESIMS *m*/*z* 350.9147 [M + H]^+^ (calcd for C_13_H_7_Cl_4_O_3_, 350.9144).

### X-ray data collection and structure refinement

Single crystals of [C_13_H_6_Cl_4_O_3_]_2_ [ethyl acetate]_0.5_ (**1**) and C_17_H_20_N_2_O_3_S_2_ (**4**) were obtained by slow diffusion of hexane into ethyl acetate solutions of the respective compounds. A suitable crystal of compound **1** was selected, mounted on a Mitegen loop using Fromblin oil, and analysed using an Xcalibur Gemini diffractometer equipped with a Ruby CCD area detector. For ulbactin B (**4**), a suitable crystal was mounted on a glass fibre with Fromblin oil and placed on a Rigaku Oxford Diffraction SuperNova diffractometer with a dual source (Cu at zero) and an AtlasS2 CCD area detector.

Crystals were kept at 150(2) K during data collection. The structures were solved using Olex2,^32^ with the ShelXT^33^ structure solution program via Direct Methods and refined using ShelXL^34^ with Least Squares minimisation.

### Crystal Data

for [C_13_H_6_Cl_4_O_3_]_2_[ethyl acetate]_0.5_ (*M* =748.01 g/mol) (CCDC 1995977): monoclinic, space group P2_1_/n (no. 14), *a* = 8.05520(7) Å, *b* = 35.3030(3) Å, *c* = 11.55129(7) Å, *β* = 92.0406(7)°, *V* = 3282.79(4) Å^3^, *Z* = 4, *T* = 150(2) K, μ(CuKα) = 6.650 mm^-1^, *Dcalc* = 1.513 g/cm^3^, 66701 reflections measured (8.058° ≤ 2Θ ≤ 156.496°), 7000 unique (*R*_int_ = 0.0496, R_sigma_ = 0.0200) which were used in all calculations. The final *R*_1_ was 0.0447 (I > 2σ(I)) and *wR*_2_ was 0.1495 (all data).

### Crystal Data

for C_17_H_20_N_2_O_3_S_2_ (*M* =364.47 g/mol) (CCDC 1995978): monoclinic, space group P2_1_ (no. 4), *a* = 6.50189(5) Å, *b* = 9.74207(6) Å, *c* = 13.67478(10) Å, *β* = 92.0045(7)°, *V* = 865.656(11) Å^3^, *Z* = 2, *T* = 150(2) K, μ(CuKα) = 2.943 mm^-1^, *Dcalc* = 1.398 g/cm^3^, 39269 reflections measured (11.154° ≤ 2Θ ≤ 147.16°), 3472 unique (*R*_int_ = 0.0724, R_sigma_ = 0.0240) which were used in all calculations. The final *R*_1_ was 0.0261 (I > 2σ(I)) and *wR*_2_ was 0.0697 (all data).

The molecule crystallised in the chiral space group P2(1) and the Flack parameter refined to - 0.001(9) (Shelx 2018) and Hooft y parameter -0.015(5) (Olex2).^35, 36^

## Acknowledgement

EM and GLC acknowledge funding from the Biotechnology and Biological Sciences Research Council (BBSRC grant BB/L021692/1) for support of metabolite discovery in *Burkholderia*. We thank Cerith Jones for technical assistance with *Burkholderia* strain revival and culture.

