## Supplementary Information for "Discovery and Metabolic Origin of 4,4′-Dihydroxy-3,3′,5,5′-Tetrachlorobenzophenone from a *Burkholderia oklahomensis* Clinical Isolate"

[1] Department of Chemistry, University of Warwick, Coventry CV4 7AL, United Kingdom
[2] School of Medical Sciences, Faculty of Medicine and Health, University of Sydney, Sydney NSW 2015, Australia
[3] Sydney Infectious Diseases Institute, University of Sydney, Sydney NSW 2015, Australia
[4] Microbiomes, Microbes and Informatics Group, Organisms and Environment Division, School of Biosciences, Cardiff University, Cardiff, UK
[5] Warwick Integrative Synthetic Biology Centre, University of Warwick, Coventry CV4 7AL, United Kingdom
[6] Department of Biochemistry and Molecular Biology, Monash University, Clayton, Australia
[7] ARC Centre of Excellence for Innovations in Peptide and Protein Science, Monash University, Clayton VIC 3168, Australia


| Contents | Page |
|---|---|
| Figure S1. $^{1}H$ NMR spectrum of 4,4′-dihydroxy-3,3′,5,5′-tetrachlorobenzophenone in DMSO-$d_6$. | S2 |
| Figure S2. $^{13}C$ NMR spectrum of 4,4′-dihydroxy-3,3′,5,5′-tetrachlorobenzophenone in DMSO-$d_6$. | S3 |
| Figure S3. COSY spectrum of 4,4′-dihydroxy-3,3′,5,5′-tetrachlorobenzophenone DMSO-$d_6$. | S4 |
| Figure S4. HSQC spectrum of 4,4′-dihydroxy-3,3′,5,5′-tetrachlorobenzophenone in DMSO-$d_6$. | S5 |
| Figure S5. HMBC spectrum of 4,4′-dihydroxy-3,3′,5,5′-tetrachlorobenzophenone in DMSO-$d_6$. | S6 |

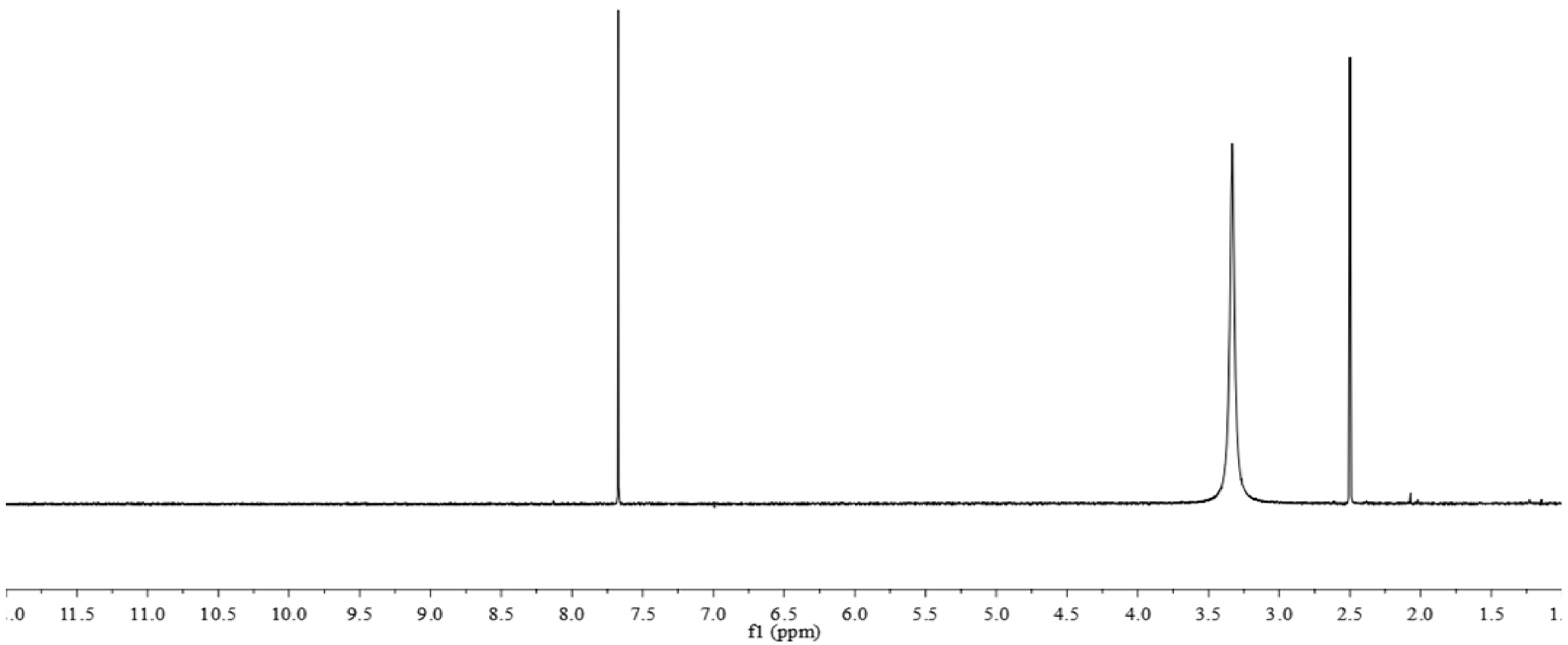


**Figure S1.** $^1$H NMR spectrum of 4,4′-dihydroxy-3,3′,5,5′-tetrachlorobenzophenone in DMSO-$d_6$.

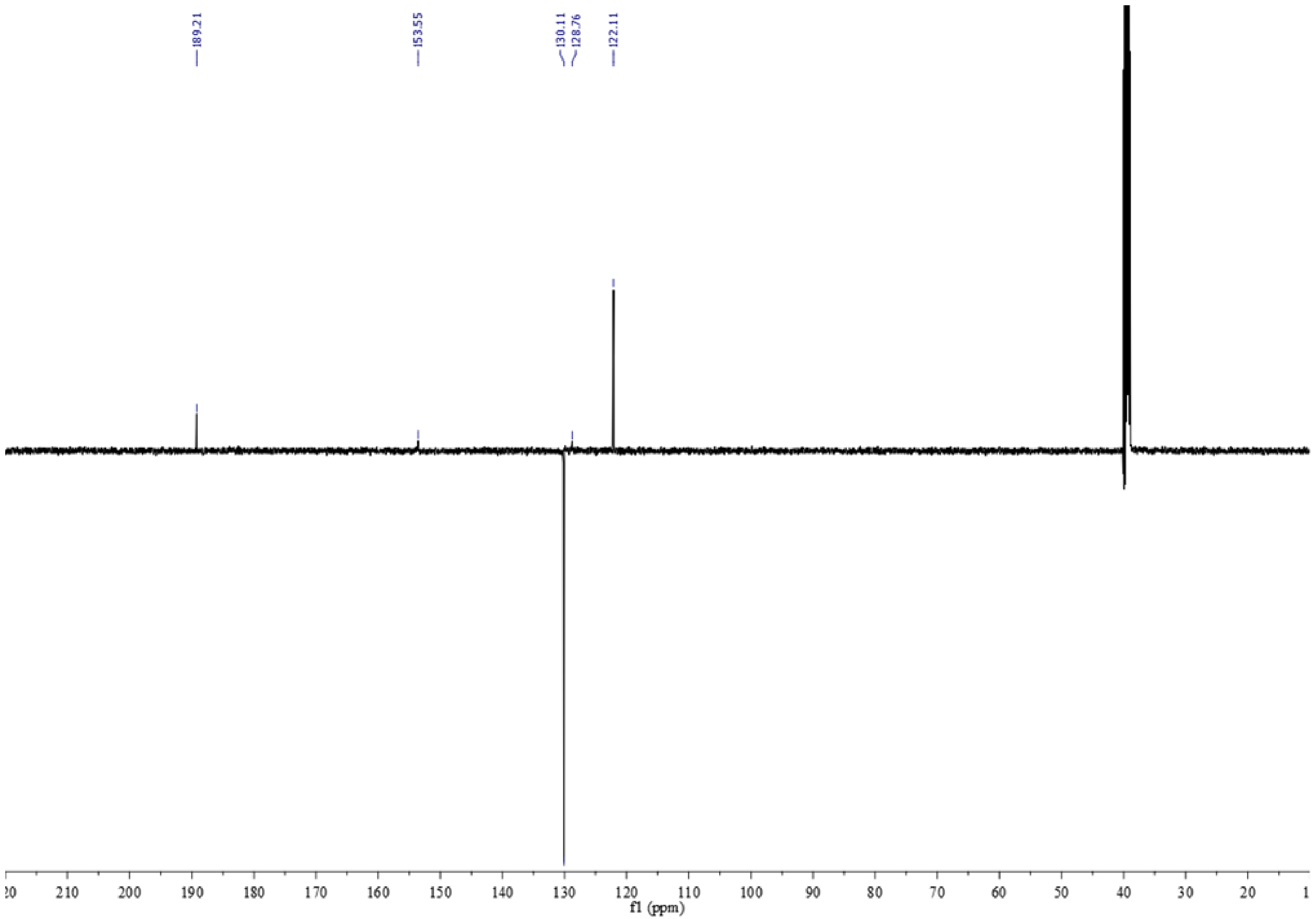


**Figure S2.** $^{13}$C NMR spectrum of 4,4′-dihydroxy-3,3′,5,5′-tetrachlorobenzophenone in DMSO-$d_6$.

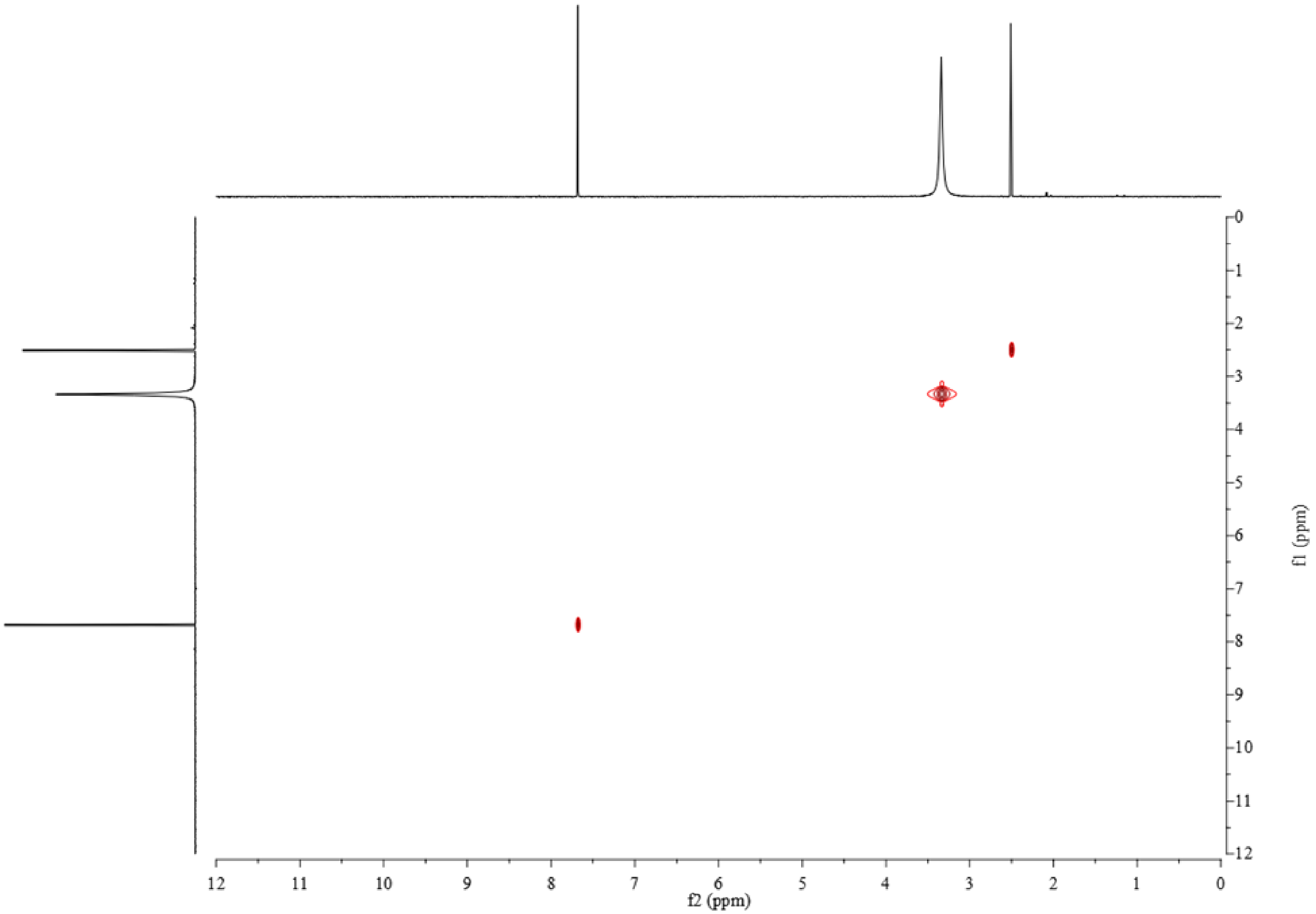

**Figure S3.** COSY spectrum of 4,4′-dihydroxy-3,3′,5,5′-tetrachlorobenzophenone in DMSO-$d_6$.

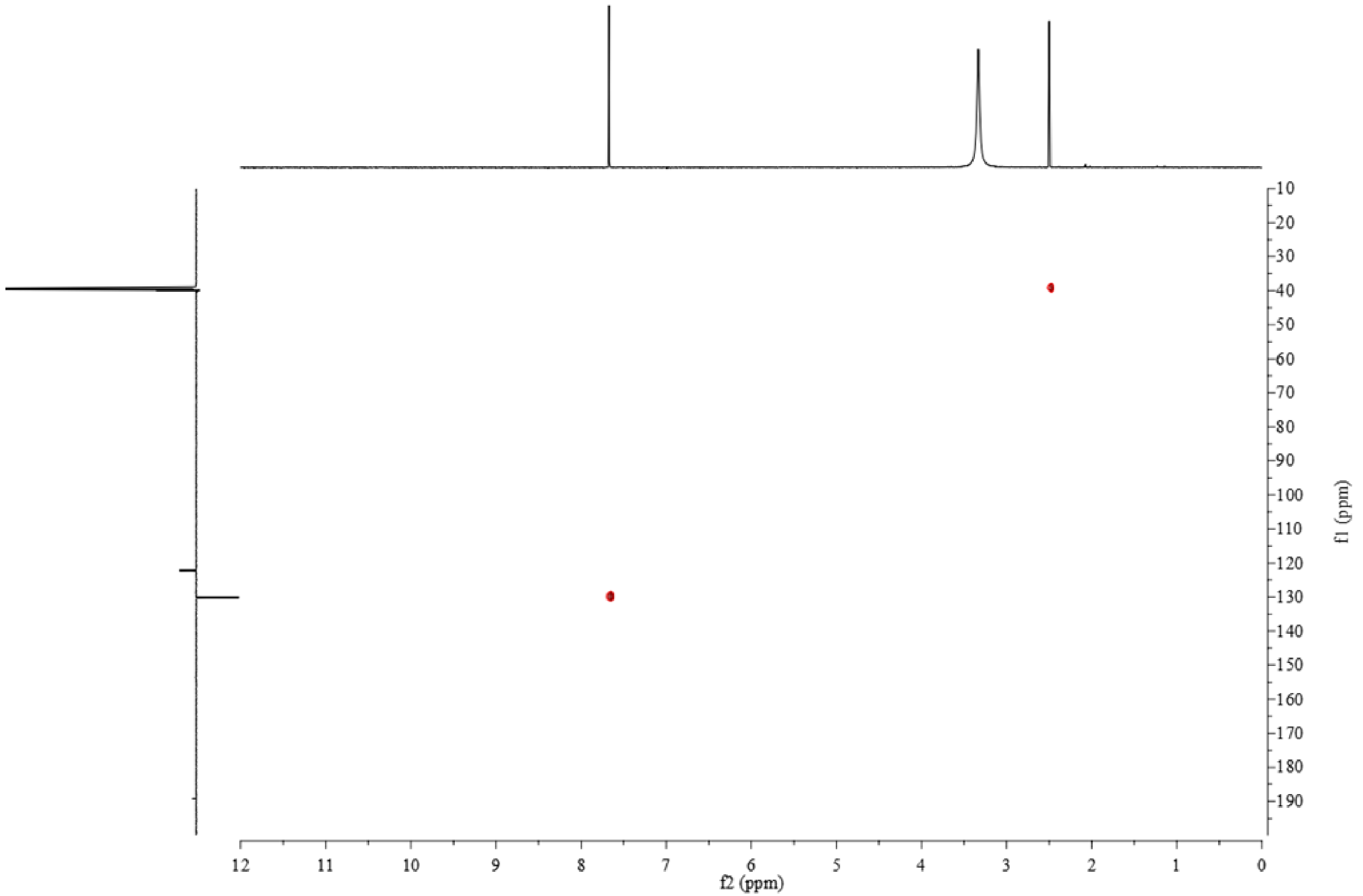


**Figure S4.** HSQC spectrum of 4,4′-dihydroxy-3,3′,5,5′-tetrachlorobenzophenone in DMSO-$d_6$.

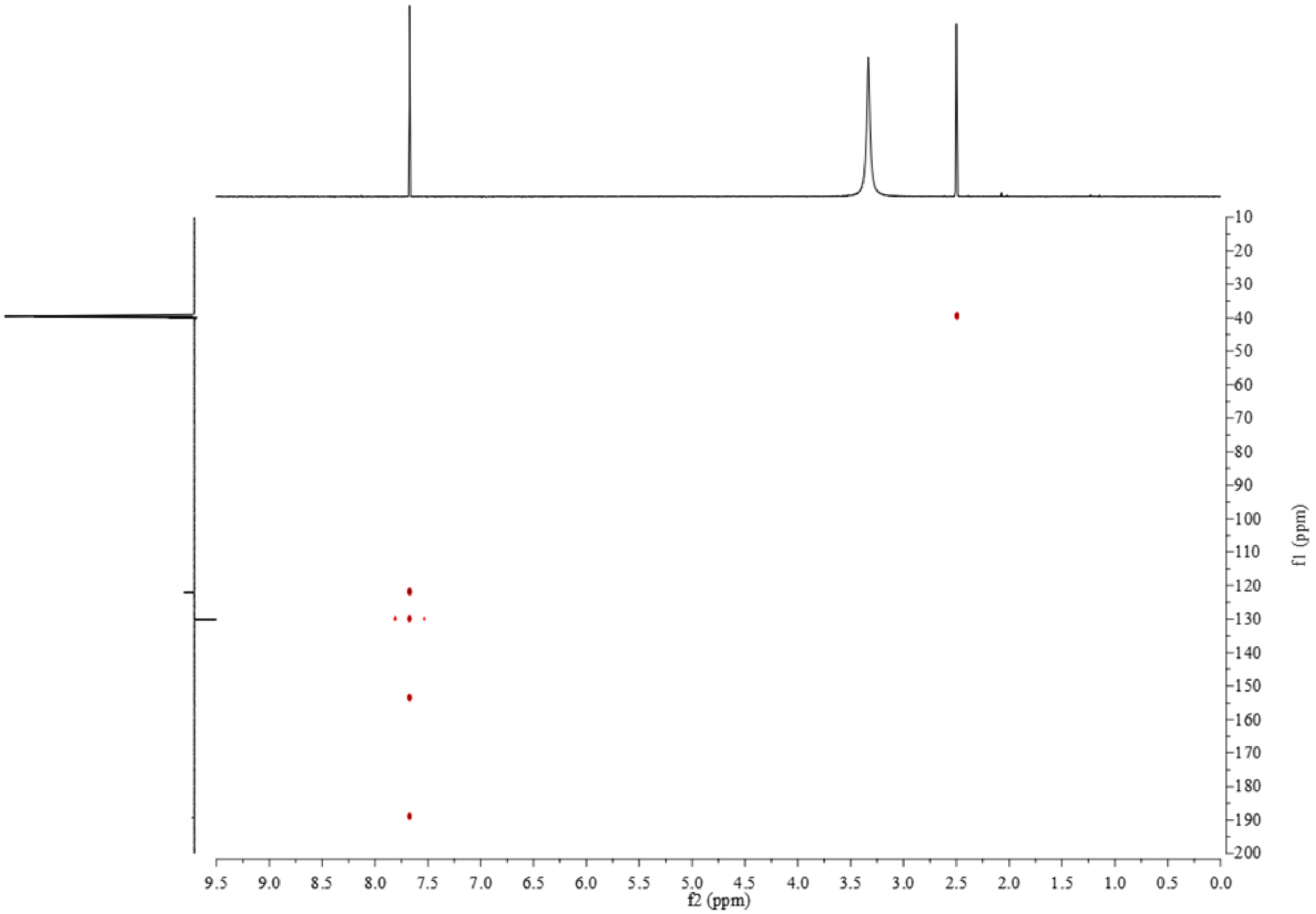


**Figure S5.** HMBC spectrum of 4,4′-dihydroxy-3,3′,5,5′-tetrachlorobenzophenone in DMSO-$d_6$.
